# The CAHRA Challenge: A Community-Wide Assessment of Cryo-EM Heterogeneous Reconstruction Algorithms

**DOI:** 10.64898/2026.09.15.751515

**Authors:** J. Ryan Feathers, Robert C. Heeter, Geoffrey Woollard, Sonya M. Hanson, Pilar Cossio, Joel Greer, Tom Burnley, Ellen D. Zhong

## Abstract

The ability of cryo-electron microscopy (cryo-EM) to interrogate the atomic structure and motion of biomolecules has motivated the development of a wide range of algorithms for heterogeneity analysis. However, evaluating and comparing these methods remains challenging because ground-truth structures are generally unknown for experimental samples. Here, we introduce the 2026 Community-Wide Assessment of Heterogeneous Reconstruction Algorithms (CAHRA), a community-wide challenge centered on three benchmark datasets that probe distinct tasks in heterogeneity analysis. These include (1) compositional heterogeneity arising from mixtures of distinct protein complexes, (2) continuous conformational variability and atomic modeling, and (3) entanglement between molecular conformation and image pose. We describe the design, construction, and validation of these datasets, as well as their use in the recently completed CAHRA Challenge.

## 1. Introduction

Single-particle cryo-electron microscopy (cryo-EM) aims to reconstruct 3-D structures of biomolecular complexes from noisy 2-D projection images. Because each projection image captures an individual molecule, a cryo-EM dataset can contain particles spanning a distribution of compositional and conformational states (1–3). Inferring both the underlying 3-D structures and their distribution from the observed set of 2-D particle images makes heterogeneous cryo-EM reconstruction a challenging inverse problem (4).

To account for structural variability, heterogeneous reconstruction algorithms (HRAs) relax the assumption that all particle images describe a single underlying structure. Instead, they implement a variety of approaches to model differences in molecular composition or conformation within a sample (5–7). These methods can identify distinct compositional states, resolve conformational ensembles relevant to molecular function, or improve consensus reconstructions by separating contaminants or damaged particles (8–13). In practice, different HRAs make different assumptions about the underlying structural distribution, which further complicates systematic benchmarking efforts (14).

Although a growing number of HRAs have been proposed, their validation remains difficult. Experimental datasets lack ground-truth information about the target structures and other particle-level variables, including structural state, image poses, and exact imaging parameters. Reconstruction accuracy can therefore be assessed only through measures such as reproducibility or agreement with published structures. Determining how faithfully reconstructed molecular motions or population estimates reflect the underlying structural distribution remains challenging, especially for dynamic biomolecular complexes.

Here we describe three benchmark datasets built for the 2026 Community-Wide Assessment of Cryo-EM Heterogeneous Reconstruction Algorithms (CAHRA) Challenge. We build upon lessons learned from prior benchmarking efforts, including CryoBench where simulated datasets were used to assess diverse forms of heterogeneity (14), and the Inaugural Flatiron Institute Cryo-EM Conformational Heterogeneity Challenge (15), which provided a pair of simulated and experimental datasets with matched conformational heterogeneity. The CAHRA challenge datasets target complementary problems in cryo-EM heterogeneity analysis: compositional heterogeneity from combining experimental datasets of different complexes, conformational heterogeneity with atomic modeling, and the entanglement of structural state with image pose. We report how each dataset was constructed, and we provide the reference labels used for assessment.

## 2. Results

### 2.1. Overview of the CAHRA Challenge

The CAHRA Challenge consists of one experimental and two simulated datasets, each organized around a distinct task in heterogeneous cryo-EM reconstruction (Figure 1a). We aimed to combine realistic sources of noise with known structural distributions and particle-level reference information. Participants were blinded to specific information and asked to recover distinct target information for each challenge (Figure 1b). During the challenge, a leaderboard was periodically updated to display the rank order of submissions. Submissions were identified by randomly generated names unless participants requested a specific identifier.

**Figure 1.**
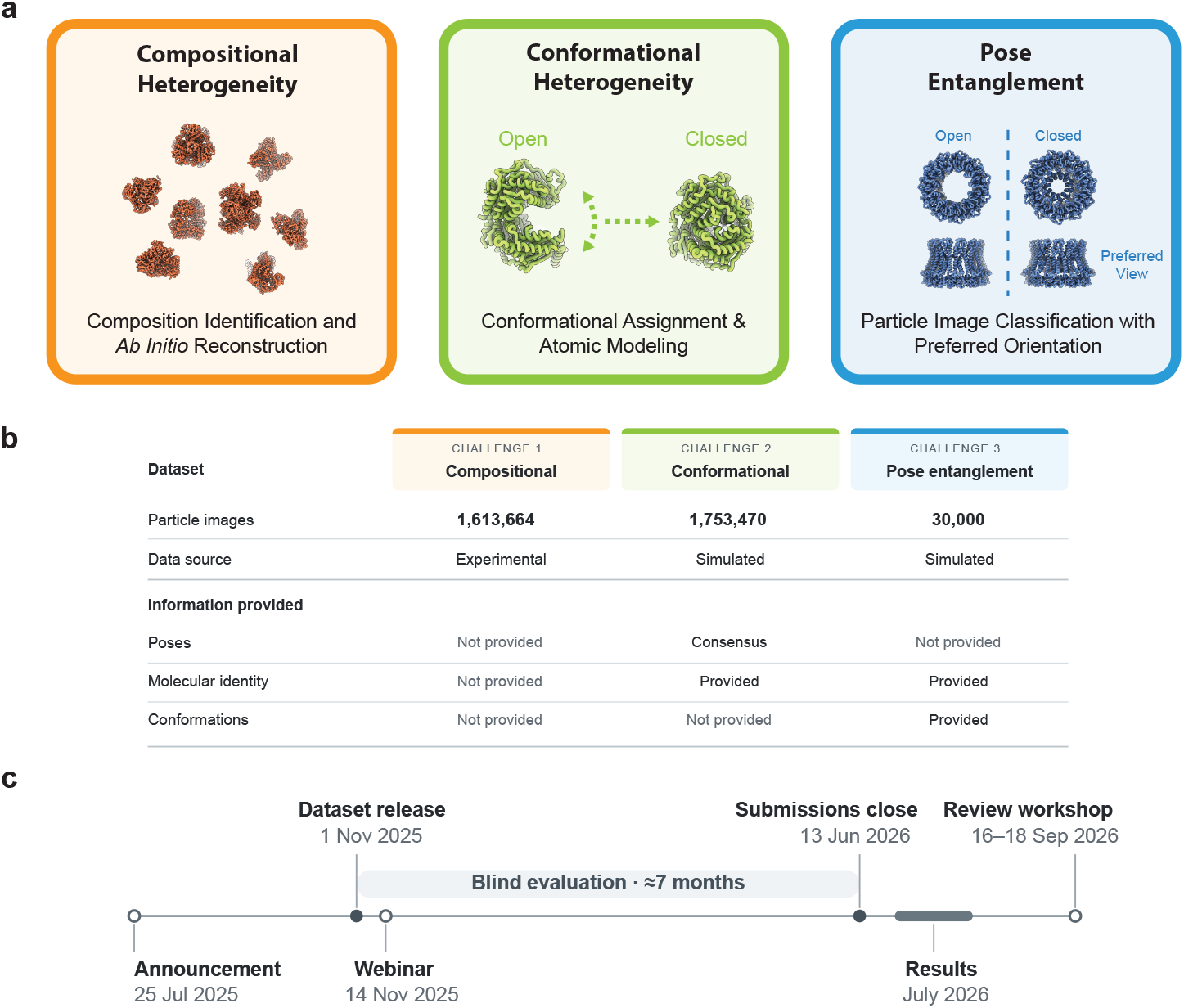
Overview of the 2026 CAHRA Challenge. **a)** The three challenges targeting distinct tasks in cryo-EM heterogeneous reconstruction and analysis. **b)** Dataset details. **c)** Competition timeline.

The challenge datasets were released and submissions opened on November 1, 2025, and participants were allowed multiple submissions during the seven-month blind evaluation period (Figure 1c). Winners of the competition were notified in July 2026. The challenge concluded with the CCP-EM CAHRA Review Workshop, held September 16-18, 2026, in Oxfordshire, United Kingdom. The workshop aimed to bring together challenge organizers, participants, and members of the cryo-EM methods development community to review the submitted approaches and discuss takeaways for the development and evaluation of HRAs.

### 2.2. Challenge 1) Compositional Heterogeneity in Complex Mixtures

Challenge 1 focused on compositional heterogeneity in a mixture of macromolecular complexes (Figure 2). Four reconstruction targets, alcohol dehydrogenase (ADH), aldolase, glutamate dehydrogenase (GDH), and pyruvate kinase (PKM2), were selected for their relatively similar sizes and combination of shared and distinct symmetries. The samples were prepared and imaged separately, but combined *in silico*, yielding ground truth particle labels for compositional state. Their similar dimensions allowed all four complexes to be accommodated within the same cropped box size, while their overlapping size and symmetry characteristics were intended to make species separation challenging. Additional imaging and processing details are provided in the Methods section.

**Figure 2.**
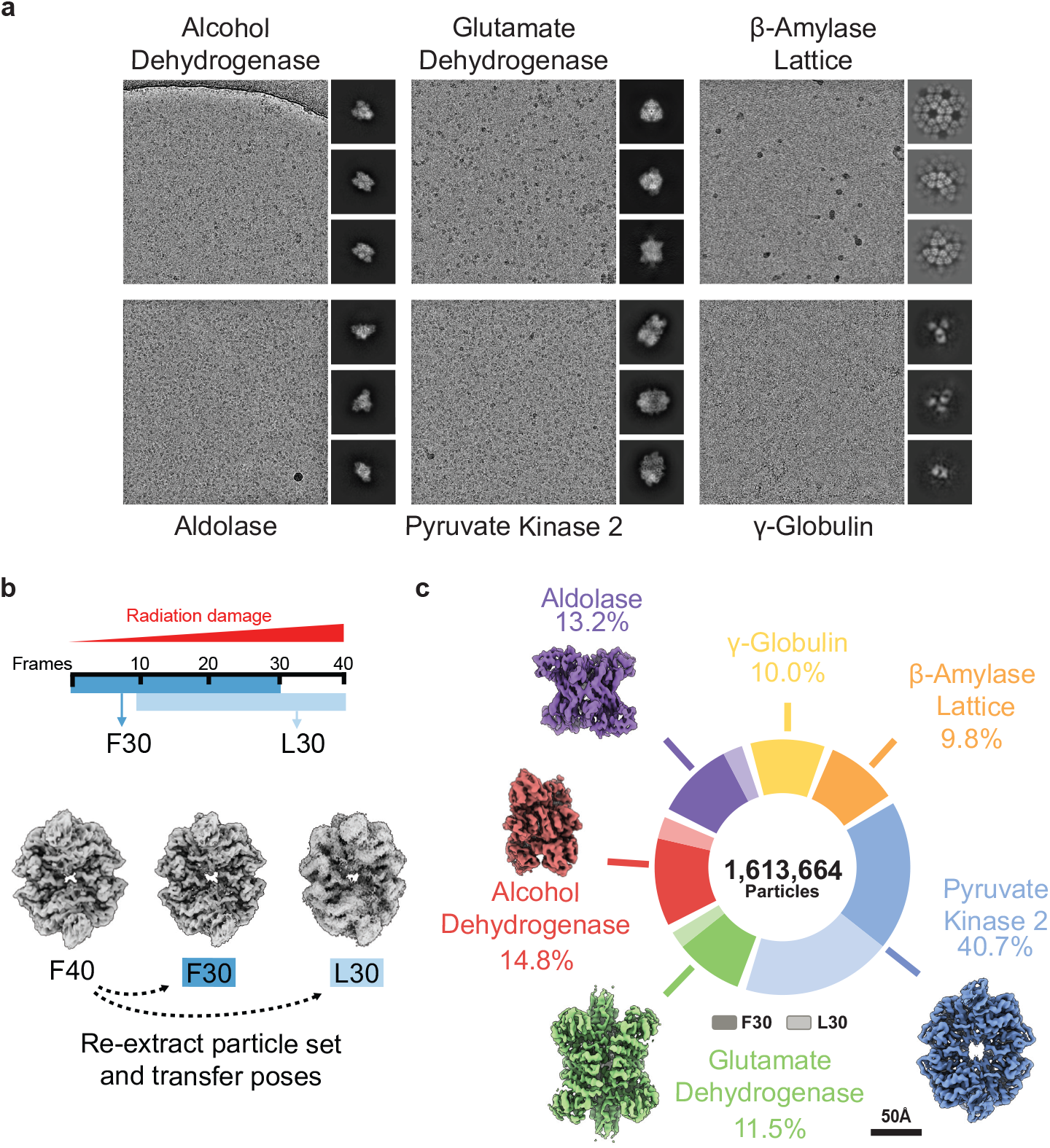
Challenge 1) Compositional Heterogeneity in Complex Mixtures. **a)** Representative experimental micrographs and 2D class averages for six purified samples. **b)** Radiation-damaged subsets were generated by re-extracting particle images summed from either the first (F) or last (L) 30 exposure frames. **c)** Final dataset composition by species and image quality. *β*-amylase and *γ*-globulin were included as decoy species.

Two additional samples, *γ*-globulin and *β*-amylase, were also imaged to serve as decoy particles with known source labels. The *γ*-globulin sample exhibited substantial structural heterogeneity, which prevented high-resolution reconstruction. Unexpectedly, the *β*-amylase preparation contained a 2-D lattice that produced high-resolution 2D class averages but lacked sufficient angular coverage for isotropic 3-D reconstruction (Figure 2a). Participants were not expected to produce 3-D reconstructions of *β*-amylase or *γ*-globulin; their inclusion was intended to make separation of the four targets more difficult. To further increase the difficulty, we subdivided particle images into higher- and lower-quality subsets by radiation damage. This was achieved by re-extracting particles from micrographs generated using either the first or last 30 frames of each movie (Figure 2b).

The final dataset comprised 1,613,664 particle images with a pixel size of 1.01 Å and a box size of 288 pixels. Patch-based defocus estimates were provided, but image poses were withheld. Participants were asked to identify and reconstruct the target complexes without prior knowledge of their identities or total number. They were instructed to produce the highest resolution reconstruction possible for each identified species using analysis methods of their choice. For each species, participants were required to submit the reconstructed volume, half-maps, and a particle metadata file identifying the particles used in their reconstruction and specifying their poses, defocus values, and per-particle class assignments. Participants were also asked to include a description of their analysis workflow.

### 2.3. Challenge 2) Conformational Heterogeneity and Atomic Modeling

Challenge 2 addressed conformational heterogeneity and the representation of a continuous motion using atomic models. The dataset was based on the transition between open and closed conformations of human presequence protease. A set of 100 reference conformations was extracted from the first 200 ns of an unsteered molecular dynamics simulation initialized from PDB entry 6XOU (16), capturing the closing of the protease from an open state (Figure 3a, b). These references defined the ground-truth conformational motion and were used to generate simulated cryo-EM particle images. Small perturbations to the ground-truth image poses were added via a 3D refinement job to reflect a realistic processing workflow and test whether approaches were robust to a small degree of pose uncertainty.

**Figure 3.**
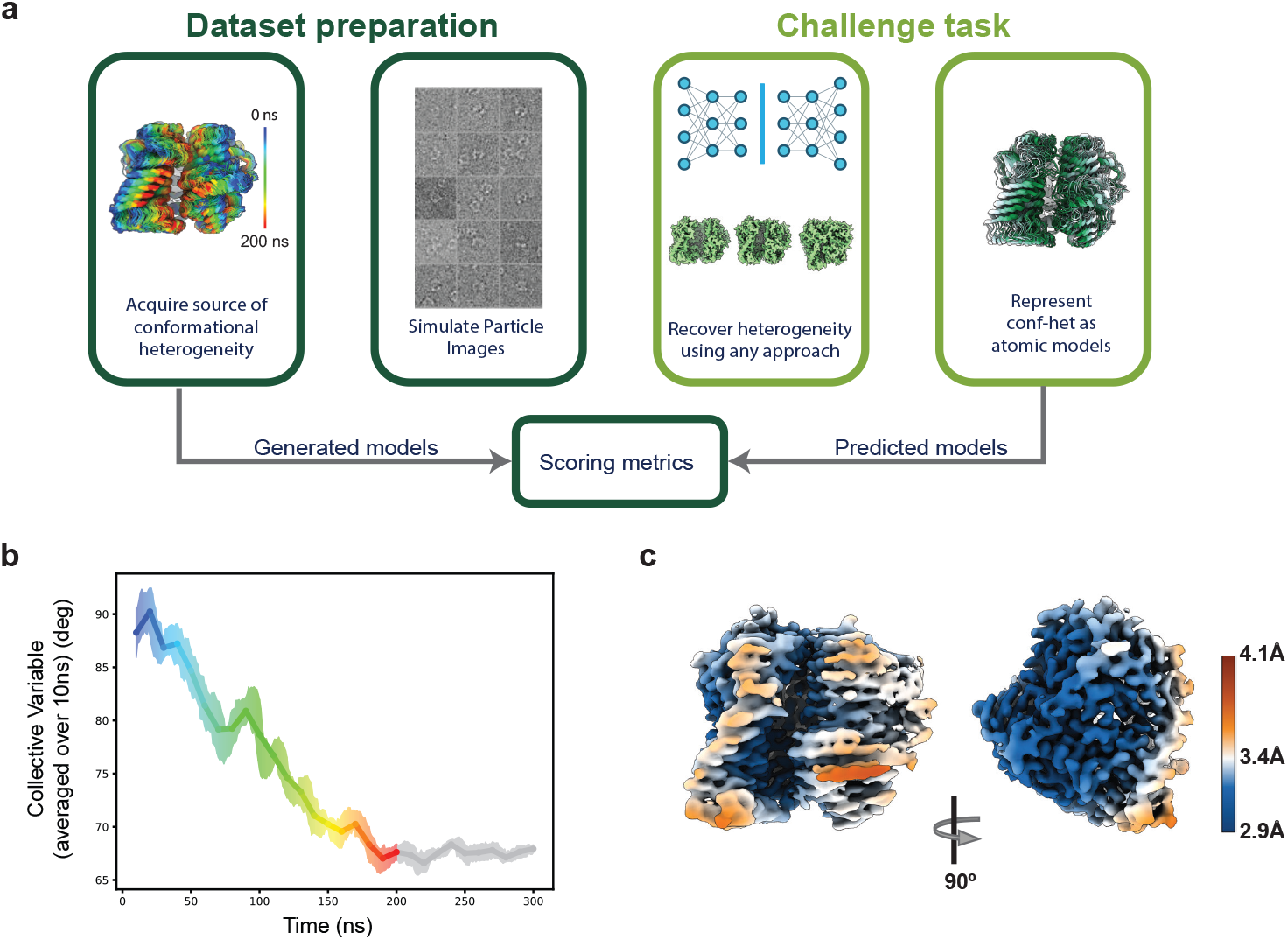
Challenge 2) Conformational Heterogeneity and Atomic Modeling. **a)** Cryo-EM particle images were simulated from snapshots along an all-atom molecular dynamics trajectory of presequence protease that captured a large-scale conformational change in the protein. **b)** A collective variable was used to parameterize the conformational change. The colors correspond to the ensemble map in (a). **c)** The local resolution of a consensus reconstruction ranged from 2.9 Å to 4.1 Å, with the lowest resolution around the active site.

The resulting dataset contained 1,753,470 particle images with a pixel size of 1.0 Å and a box size of 256 pixels. Consensus reconstruction poses, CTF parameters including per-particle defocus values, and the FASTA sequence of the protein were provided. Participants were asked to determine a set of 10-100 atomic models that collectively represent the conformational variability present in the particle stack. The option of submitting as few as 10 or as many as 100 models was intended to accommodate both manual workflows and automated pipelines.

Requiring submissions in the form of atomic models served two purposes. First, comparing sets of atomic structures substantially simplifies evaluation, as metrics to compare conformationally varying volumes are not established or standardized. Second, we aimed to evaluate recent HRA methods that combine atomic model priors with reconstruction and/or new approaches for atomic modeling of heterogeneous cryo-EM density maps. The challenge therefore emphasized both recovery of the underlying molecular motion and the accuracy of the inferred atomic models.

### 2.4. Challenge 3) Pose Entanglement

Challenge 3 was designed to assess how non-uniform pose distributions affect the recovery of conformational heterogeneity. Experimental single-particle cryo-EM datasets rarely contain uniformly distributed particle views (17). Strongly anisotropic angular sampling can produce direction dependent resolution and elongation artifacts in undersampled orientations (18). When a conformational change occurs primarily along the same axis, these reconstruction artifacts may obscure the structural difference or become entangled with estimates of conformational state.

Simulated cryo-EM images were generated from open and closed structures of calcium homeostasis modulator 2 (CALHM2) (19). CALHM2 is a ring-like channel with cyclic symmetry that undergoes a concentric conformational change around its Z axis. Projections perpendicular to the Z axis lack obvious visual differences (Figure 4). The pose and state distributions were sampled independently, such that both the open and closed states experienced the same preferred orientation bias. Participants were asked to recover the underlying structural states (both global fraction and per-particle assignments) and per-particle pose. Only per-particle CTF and open/closed volume templates were provided. Particle images were also generated with a range of noise levels to study the effect of noise level on pose and state estimation.

**Figure 4.**
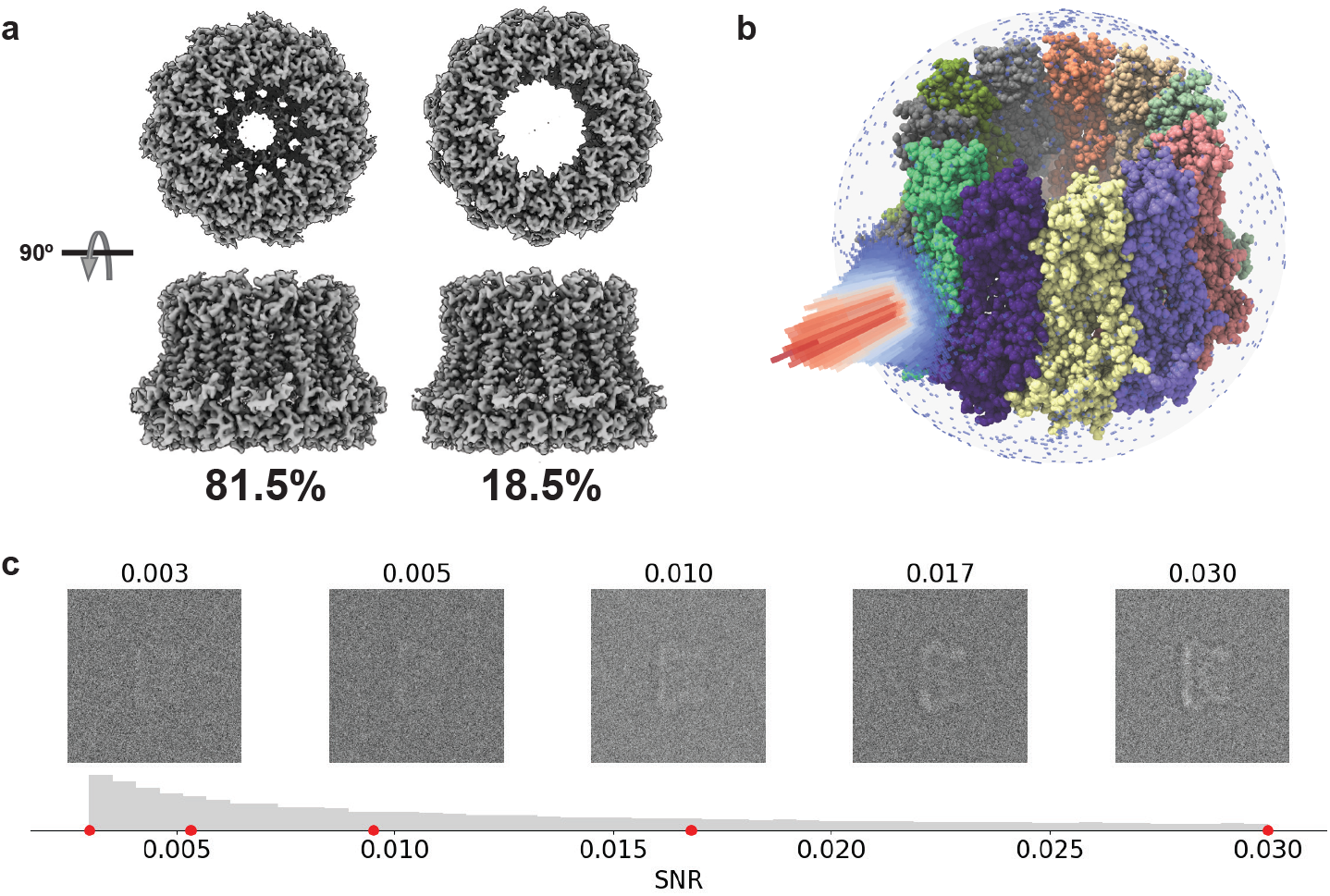
Challenge 3) Pose Entanglement. **a)** Cryo-EM images were simulated with a ratio of 81.5% closed-state to 18.5% open-state particles. **b)** Images were generated with a strong bias towards a side view of the complex, obscuring the variable region. **c)** Particle images were corrupted with noise levels sampled from a log-uniform distribution.

This challenge was intended to determine whether heterogeneous reconstruction methods could accurately quantify the fraction of open and closed states, and to what extent state prediction was influenced by preferred orientation.

## 3. Outlook

This manuscript describes the design and construction of the three CAHRA benchmark datasets that address distinct limitations in the current evaluation of heterogeneous reconstruction algorithms. Following the CCP-EM CAHRA Review Workshop, we will provide an update reporting the full competition results and our analysis.

In compositional heterogeneity, the ability to separate multiple molecular species from experimental images has been demonstrated in increasingly complex samples. Nevertheless, such analyses often remain computationally intensive and depend on iterative parameter tuning, manual inspection, and decisions made by experienced users. Challenge 1 is therefore intended not only to test whether four target complexes and state populations could be recovered, but also to provide a standardized setting for developing workflows that perform heterogeneous *ab initio* reconstruction more efficiently, reproducibly, and with less user intervention.

Continuous conformational heterogeneity presents a different challenge for both reconstruction and evaluation. Challenge 2 provides a large simulated dataset of a single protein with a defined conformational motion and realistic image-formation effects. Unlike prior conformational heterogeneity benchmarking efforts (15), by requiring submissions as atomic models, Challenge 2 provides a common representation for quantitative evaluation while testing the end-to-end recovery of conformational variability from particle images to atomic structures. More broadly, benchmarks that connect heterogeneous reconstruction to atomic modeling will be important for assessing whether recovered structural ensembles faithfully capture molecular motions at biologically relevant scales.

Challenge 3 highlights the importance of evaluating structural heterogeneity together with other latent variables in the cryo-EM imaging process. Particle pose is typically estimated jointly with, or prior to, structural variability, and errors in pose estimation can become coupled to estimates of conformational state. Preferred orientation provides a particularly challenging setting because the most frequently observed views may contain limited information about the underlying structural differences. By providing ground-truth poses and states for evaluation, Challenge 3 enables direct measurement of this coupling. Similar benchmarks could examine interactions between heterogeneity and other sources of variation in experimental data, helping determine when apparent structural variability reflects the underlying molecular ensemble rather than errors or biases introduced during reconstruction.

Together, the three datasets and their associated ground-truth information provide a resource for the continued development of heterogeneous reconstruction methods. Although workflows and best practices for handling some types of heterogeneity exist, substantial work remains to establish the reproducibility and robustness of these workflows, especially to the complexities of experimental data. Developing these approaches will expand the capabilities of cryo-EM to investigate increasingly complex samples, which will become increasingly important in applications such as time-resolved cryo-EM, *in situ* single-particle analysis, and cryo-electron tomography, where resolving heterogeneity is central to the long-term goal of visual proteomics.

## 4. Methods

### 4.1. Challenge 1) Compositional Heterogeneity

The Challenge 1 dataset was assembled from cryo-EM particle images collected independently from six purified protein samples: yeast alcohol dehydrogenase, rabbit aldolase, bovine glutamate dehydrogenase, human pyruvate kinase 2, sweet potato *β*-amylase, and bovine *γ*-globulin. All samples were imaged during a single 24-hour session on a Cs-corrected Titan Krios G3 equipped with a Falcon 4i detector and Selectris energy filter, using a pixel size of 1.01 Å/pixel and a total exposure of approximately 50 *e*^−^/Å2. Movies were motion corrected and dose weighted, followed by CTF estimation and sample-specific particle picking and classification. Single particle analysis was performed on each sample separately to verify that the challenge reconstruction targets contained a subset of particles capable of achieving 4 Å resolution. The volumes shown in Figure 2 were reconstructed by homogeneous refinement using selected subsets of 145,074 particles for ADH, 145,633 for aldolase, 99,777 for GDH, and 235,178 for PKM2. These reconstructions verified that the source data for each target contained sufficient images to achieve roughly 4 Å resolution or better. The subsets do not necessarily represent optimal particle selections, and higher resolution reconstructions may be achievable with additional analysis.

Following this validation, unfiltered particle coordinates from the upstream particle picking step were selected as the base population for each molecular species. *β*-amylase was initially intended to serve as a fifth reconstruction target; however, the appearance of a 2D lattice likely resulting from a degradation dominated the micrographs. Particle images from the six independently processed datasets were then combined into a single challenge dataset. Testing and tuning the difficulty of mixed class *ab initio* reconstruction was then performed using common workflows in RELION and CryoSPARC (8, 9).

To increase the difficulty of this challenge, the exposure was truncated to 30 fractions by excluding either the first 10 frames (L30) or the last 10 frames (F30) during motion correction. The proportion of randomly selected radiation-damaged particles was then increased or decreased for each species to tune the difficulty of separation. The percentages of L30 particles included for each species were as follows: PKM2 (50%), ADH (20%), aldolase (20%), and GDH (20%). No L30 particles were included for *β*-amylase or *γ*-globulin. To obscure their source identities, particles were reordered and regrouped by similar defocus values, sequential micrograph names were assigned, and small perturbations were added to the CTF angle. The final STAR file provided to participants contained the particle images and processing metadata but no sample identities or reference labels.

### 4.2. Challenge 2) Conformational Heterogeneity

The Challenge 2 dataset was generated synthetically from an atomistic molecular dynamics simulation of human presequence protease initialized from the open conformation (PDB 6XOU). Missing residues were modeled and the structure refined before minimization, equilibration, and a 1 ??s molecular dynamics simulation. A collective variable describing opening and closing of the molecule was used to characterize the trajectory, and 100 reference conformations were selected from the first 200 ns to span the observed conformational range (Figure 3b). These structures were used with Roodmus and Parakeet to simulate 6,668 cryo-EM micrographs with varying defocus and representative microscope parameters, including a 300 kV accelerating voltage, 2.7 mm spherical aberration, 45 *e*^−^/Å2 total fluence, and a pixel size of 1.0 Å (20, 21). Reference conformations were uniformly sampled in the creation of these micrographs. Particles were extracted in RELION with a 256-pixel box size after removing particles near micrograph edges. A consensus 3D refinement was then performed to introduce small perturbations to the ground-truth image poses and translations. The resulting particle stack, refined poses, per-particle defocus estimates, detector modulation transfer function, and protein sequence were provided to challenge participants, while the reference conformations and associated ground-truth metadata were withheld.

### 4.3. Challenge 3) Pose Entanglement

The Challenge 3 dataset consists of synthetic particle images that were generated using CryoJAX (version 5) from open and closed atomic structures, 6UIV and 6UIW respectively, modified to have an identical number of atoms (i.e. no missing/extra residue discrepancies or ligand differences) and each equilibrated in a short MD simulation, at a fixed population ratio of 0.185:0.815. Particle orientations were sampled from a strongly non-uniform distribution centered on a side view, with 3% of particles assigned uniformly distributed orientations to provide limited angular coverage outside the dominant view. Images were generated directly using CryoJAX (22) from non-hydrogen atomic coordinates using Gaussian mixture projection and simulated with a 300 keV accelerating voltage, 2.7 mm spherical aberration, 0.1 amplitude contrast, a defocus range of 1–2 ??m, a pixel size of 1.2 Å, and a 256-pixel box size. Gaussian white noise was added with particle-specific signal scaling sampled over a broad range to introduce variation in image quality. The resulting dataset contained 30,000 particles with known conformational states, poses, and imaging parameters. State, pose and noise level were withheld from participants during the challenge.

## 5. Data Availability

The CAHRA Challenge datasets and associated ground-truth particle labels, reference structures, and evaluation metadata are publicly available in the Electron Microscopy Public Image Archive under accession numbers EMPIAR-13973 (compositional heterogeneity), EMPIAR-14006 (conformational heterogeneity), and EMPIAR-13991 (pose entanglement). Additional data can be made available upon request.

## 6. Author Contributions

All authors conceived the study. P.C., S.M.H., T.B., and E.D.Z. supervised the study. J.R.F. collected experimental cryo-EM data. J.R.F. and R.C.H. prepared the compositional heterogeneity dataset. J.G. prepared the conformational heterogeneity dataset. G.W. prepared the pose entanglement dataset with assistance from S.M.H. J.R.F. wrote the initial version of the manuscript. All authors wrote the final version of the manuscript.

## 7. Acknowledgments

The authors thank Jonathan Bouvette, George Ghanim, and Rishwanth Raghu for assistance with data collection and preliminary testing; Frederick Hughson for helpful discussions regarding the compositional heterogeneity dataset; and Miro A. Astore and Luke Evans for helpful discussions regarding the pose entanglement dataset. The authors are grateful to Renaissance Philanthropy for sponsoring competition prizes.

J.R.F., R.C.H., and E.D.Z. acknowledge the use of computing resources at Princeton Research Computing, a consortium of groups led by the Princeton Institute for Computational Science and Engineering (PICSciE) and Office of Information Technology’s Research Computing. J.R.F., R.C.H., and E.D.Z. are supported by the Chan Zuckerberg Initiative DAF (grant number 2025-358484), an advised fund of Silicon Valley Community Foundation, the National Institutes of Health (grant number DP2GM164606), and the AI2050 program at Schmidt Sciences (grant number G-25-69788). E.D.Z. acknowledges funding support from the Princeton Catalysis Initiative, Princeton School of Engineering and Applied Sciences, Janssen Pharmaceuticals, and Generate Biomedicines. The Flatiron Institute is a division of the Simons Foundation. This project made use of time on HPC granted via the UK High-End Computing Consortium for Biomolecular Simulation, HECBioSim (http://hecbiosim.ac.uk), supported by EPSRC (grant number EP/X035603/1). J.G. and T.B. are supported by UK Research and Innovation through Medical Research Council (grant number MR/V000403/1, grant number UKRI2367) and through funding for the project entitled ‘2024 CoSeC CCP Bridging: Digital Research Infrastructure for Integrative Molecular Biology (DRIIMB)’ awarded by the UKRI Science and Technology Facilities Council 2024-2026. G.W. is supported by an NSERC Canada Graduate Scholarship – Doctorate, and was supported by an NSERC Michael Smith Foreign Study Supplement for this research while visiting the Flatiron Institute. These funders had no role in study design, data collection and analysis, decision to publish, or preparation of this manuscript.

