## Extended Methods for "The CAHRA Challenge: A Community-Wide Assessment of Cryo-EM Heterogeneous Reconstruction Algorithms"

### Supplementary Information for: The CAHRA Challenge: A Community-Wide Assessment of Cryo-EM Heterogeneous Reconstruction Algorithms

**J. Ryan Feathers[1,2], Robert C. Heeter[3], Geoffrey Woollard[4,5,6], Sonya M. Hanson[5,6], Pilar Cossio[5,6], Joel Greer[7], Tom Burnley[7] and Ellen D. Zhong[1,3]**

[1]Department of Computer Science, Princeton University, Princeton, NJ, USA, [2]Department of Molecular Biology, Princeton University, Princeton, NJ, USA, [3]Omenn-Darling Bioengineering Institute, Princeton University, Princeton, NJ, USA, [4]Department of Computer Science, University of British Columbia, Vancouver, BC, Canada, [5]Center for Computational Biology, Flatiron Institute, New York, NY, USA, [6]Center for Computational Mathematics, Flatiron Institute, New York, NY, USA, [7]Scientific Computing Department, Science and Technology Facilities Council, Harwell, UK

#### Abstract

#### S1. Extended Methods

##### S1.1. Challenge 1 dataset preparation

###### *S1.1.1. Sample preparation*

The compositional heterogeneity challenge dataset was created by combining extracted particle images from five purified samples of commercially available proteins and one recombinantly expressed and purified protein. Yeast alcohol dehydrogenase (Sigma-Aldrich, A8656), rabbit aldolase (Sigma-Aldrich, A2714), bovine glutamate dehydrogenase (Sigma-Aldrich, G7882), and sweet potato $\beta$-amylase (Sigma-Aldrich, A7005) were resuspended and buffer exchanged into 25 mM HEPES, pH 7.5, 150 mM NaCl, and 1 mM TCEP using Zeba desalting spin columns. Samples were then filtered using 0.22-$\mu$m spin filters before plunge freezing. Bovine $\gamma$-globulin was purified from gel-filtration standards (Bio-Rad, 1511901) by size-exclusion chromatography using 25 mM HEPES, pH 7.5, and 150 mM NaCl as the running buffer. Human PKM2 was recombinantly expressed in *Escherichia coli* and purified by nickel-affinity and size-exclusion chromatography. pET-28a-hPKM2 was a gift from Lewis Cantley & Matthew Vander Heiden (Addgene plasmid # 44242 ; http://n2t.net/addgene:44242 ; RRID:Addgene_44242)(1). The PKM2 size-exclusion chromatography running buffer additionally contained 5 mM $MgCl_2$ and 1 mM fructose 1,6-bisphosphate (Neta Scientific, AST-D81856).

Samples were diluted to their final working concentrations before grid preparation: 3.0 mg/mL alcohol dehydrogenase, 1.5 mg/mL aldolase, 1.0 mg/mL glutamate dehydrogenase, 1.0 mg/mL PKM2, 0.5 mg/mL $\beta$-amylase, and 0.5 mg/mL $\gamma$-globulin. Quantifoil 1.2/1.3 grids were glow discharged for 60 seconds at 15 mA and 0.40 mBar. For each protein complex, a 3-$\mu$L volume sample was applied to a grid, blotted for 2–4 s using a blot force of 0, and plunge frozen using a Thermo Fisher Scientific Vitrobot.

###### *S1.1.2. Cryo-EM data collection*

Grids prepared from each sample were screened before data collection. Data for all six samples were collected during a single 24-hour imaging session to minimize differences in optical aberrations resulting from changes to the microscope optics. In total, 5,628 movies were recorded on a Cs-corrected Titan Krios G3 microscope equipped with a Falcon 4i detector and a Selectris energy filter operated with a 10-eV slit width. Movies were acquired over a nominal defocus range of −1.0 to −2.0 $\mu$m at a nominal magnification of 105,000×, corresponding to a pixel size of 1.01 Å/pixel, with a total exposure of approximately 50 $e^{-}/\text{Å}^{2}$. Data were saved in electron-event representation (EER) format with 945 raw frames per movie.

###### *S1.1.3. Cryo-EM data analysis*

Cryo-EM data for 6 purified protein samples were processed using RELION 5 and cryoSPARC(2, 3). RELION 5 implementation of motioncorr2 was used for motion correction and dose weighting with an EER raw frame grouping of 23 resulting in 40 fractions. Motion corrected summed micrographs were imported into cryoSPARC and defocus values were estimated using patch-CTF. CryoSPARC blob picker was used to pick particles from 282 Aldolase micrographs and 1,049 PKM2 micrographs. Alcohol dehydrogenase particles were picked from 870 micrographs using 2D template picking with class averages obtained from performing blob picking and 2D classification on a subset of the micrographs. Glutamate dehydrogenase particles were picked from 1,481 micrographs using 2D template picking with 25 2D templates simulated from a volume obtained during screening. Unexpectedly, beta-amylase appeared to form a 2D crystal lattice at all concentrations screened. Blob picking and 2D classification revealed a lattice structure with ring like features. Particles were picked from 238 micrographs using a 3D template generated from the crystal symmetry model of 1FA2. SDS-PAGE later revealed high concentration of a monomer of lower molecular weight than the beta-amylase monomer. Particle images obtained from 2D classification of this sample were included but this sample likely lacks the Fourier coverage necessary to produce an isotropic 3D reconstruction. Bovine gamma-globulin was chosen to include as a sample which is too heterogeneous to produce a high resolution average. Low resolution 2D templates were generated by manually picking performing 2D classification on a subset of the 409 micrographs. 2D template picking and class averaging were performed before selecting a subset of the data with strong low resolution features for re-extraction and inclusion in the challenge dataset.

###### *S1.1.4. Challenge 1 dataset composition*

STAR files for each set of particle images containing defocus values were combined. Particles were re-ordered and regrouped with particles of similar defocus values and sequentially numbered micrograph names were added. A small amount of jitter was added to the CTF angle to further anonymize the reference groupings. Challenge participants were provided with a STAR file stripped of all reference labels.

##### S1.2. Challenge 2 dataset generation

The conformational heterogeneity challenge utilizes an entirely synthetic dataset. As per section S1.2.1, an atomic model was sourced from the Protein Data Bank (PDB) (4), missing residues were built and the structure was refined to get a starting structure for a molecular dynamics (MD) simulation. This structure was minimized and equilibrated before it was used to run an atomistic MD simulation (section S1.2.2). The conformational change was analyzed, making use of a collective variable (CV). A subset of conformations was extracted to form an ensemble of ground truth atomic models as reported in section S1.2.3.

The ground truth ensemble was used to simulate a set of synthetic micrographs populated by the selected conformations. Ground truth metadata was to create a RELION 3.1 STAR file (5) which was utilized to extract the synthetic particle set. The filtered particle stack(s) and the 3D-refined particles STAR file were provided to entrants along with the detector modulation transfer function and the FASTA sequence of the protein.

##### *S1.2.1. Model building and refinement*

The atomic model with PDB ID 6xou (6) was downloaded from the PDB as an initial starting structure. This corresponds to human presequence protease in the "open" conformation as determined by electron cryo-microscopy. The corresponding density map and half-maps deposited in the EMDB (7) entry EMD-22280 were utilized for building missing residues and refining the model. Building missing residues was done using COOT 0.9.8.4 (8).

Firstly, despite poor Coulomb potential in the region, the N-terminal histidine tag reported to be present in experiment was manually built following the sequence reported in the wwPDB EM validation report.

Residues 806-847 were not built in 6xou, where these residue sequence numbers are those corresponding to the UniProt (9) sequence Q5JRX3. An AlphaFold2 (10) prediction for UniProt sequence Q5JRX3 was downloaded and overlaid in COOT using a least squares (LSQ) fit. The missing residues were transplanted into the atomic model from the aligned AlphaFold2 prediction. Conflicting residues identified using the validation report were then mutated to those corresponding to UniProt ID Q5JRX3.

This model was then refined using Refmac-Servalcat (11) for 20 epochs. Model validation was run on the refined atomic model and used to inform a round of manual atomic refinement. Further Refmac-Servalcat refinement with default parameters did not further improve validation statistics compared to the manually refined model, so this was taken forward.

##### *S1.2.2. Molecular dynamics simulation*

The provenance data of the preparation, equilibration and minimization steps to prepare the atomic model for "production runs" of MD was captured using the aiida-gromacs (12) and aiida-amber tools (13).

Firstly, pdb2pqr (14) was used to generate a PQR file from the atomic model using an AMBER forcefield at 7.7 pH. Molecular topology and coordinate files were then generated by tleap via AmberTools (15).

Initial minimization of the system was carried out in 3 steps using AMBER's sander engine (16). The first of these was an unrestrained energy minimization, the second step utilized positional restraints on the solute leaving only the waters free to move and the third step was a 10 ps constant-volume MD heating (equilibration) run in steps of 1 fs. The resulting AMBER parameter-topology and restart-coordinate files were then converted into GROMACS (17) format using the ParmEd package (18). Note that the residue sequence numbers are altered by AMBER so that in downstream PDB files the N-terminal of the histidine tag starts from an index of 1.

The GROMACS preprocessor was used to create GROMACS portable binary run input files for another 3 steps of energy-minimization and equilibration which were run via GROMACS. The first (energy minimization) step used a loose convergence criterion of 1000 kJ/mol/nm to resolve bad clashes. The second step was a constant-volume, constant-temperature (NVT) equilibration for 100 ps in 2 fs steps at a temperature of 300 K. This was followed by a constant-pressure, constant-temperature (NPT) equilibration for another 100 ps, again in 2 fs steps. The NPT equilibration was followed by 1 μ s of production (NPT) MD simulations in 100 ns steps.

##### *S1.2.3. Selection of reference conformations*

The MD trajectories were joined using the GROMACS trajectory concatenation tool before the GROMACS trajectory conversion tool was used to center frames within the period boundary conditions, before rotationally and translationally fitting to first frame, which was used as the reference structure.

The ground truth ensemble was curated to explore as large a conformational change as possible with simplicity being preferred over complexity in the course of this change. A collective variable (CV) was used to parameterize the opening/closing of the PreP molecule. This CV was defined as the angle between center of masses of the D1 subdomain, the linker between the two domains and the D4 subdomain, following that used in (6).

It was found that in the course of the first 200 ns of the MD simulation, the CV decreased from 87°to 67°, thereafter oscillating about this value. Using MDAnalysis (19)(20)(21)(22), a principal component analysis (PCA) of the first 200 ns of the trajectory showed that 81.5% of the variance was explained by the first principal component (PC), with the next 10.5% of the variance being cumulatively explained by the next 9 PCs. Whilst the MD simulation was unbiased and atomistic, the cosine content of the first principal component was 0.93, clearly showing that this motion was not random diffusion and that it may be unwise to make inferences about real-world protein function from this MD simulation. Sampling the ground truth ensemble from this conformational change was considered sufficient to create a benchmarking dataset.

The ground truth ensemble was constructed from 100 conformations extracted from the first 200 ns of the MD simulation. These conformations were sampled uniformly across the range of the CV after removing outliers with high CV angles (frames with base-0 indices 77, 107) which would have caused duplication of selected frames.

##### *S1.2.4. Synthetic dataset generation and curation*

The ground truth ensemble was provided to Roodmus (23) to generate a set of 6668 synthetic micrographs using Parakeet as the image simulation backend (24). The dataset was generated in two parts. The first 3334 micrographs having a defocus sampled from an normal distribution with a mean of 2µ̃m and standard deviation of 0.5 µm. The second 3334 micrographs had defoci sampled Other micrograph simulation parameters include from a set of normal distributions with means of 1 µm to 1.9 µm in 0.1 µm intervals and standard deviations of 0.05 µm. A full list of configuration parameters are noted in the deposited bash scripts, with any not noted using default values. The most important parameters were a total electron fluence of 45 Åper micrograph, an electron beam energy of 300 kV, a spherical aberration of 2.7 mm and a chromatic aberration of 2.7 mm and a pixel size of 1.0 Å. The synthetic micrographs were simulated as "single exposures" rather than movies.

Roodmus was used to create a RELION 3.1 STAR file with per-particle defoci. Particles at the edge of micrographs were filtered out of the dataset but particles which overlapped in the x-y plane were not. RELION (25) was used to extract the particles with a box size of 256 pixels with no down-sampling. A consensus 3D-refinement was then run to provide a perturbation to the ground truth positions and orientations. This utilized Blush regularization (26), a mask of 120 Å, an initial low pass filter of 10 Åand an angular sampling interval of 1.8°and an offset search range of 2 pixels in 1 pixel steps to provide relatively small perturbations. The resulting particle translations and orientations were released for use in the conformational heterogeneity challenge.

##### S1.3. Challenge 3 dataset generation

The pose entanglement challenge was designed to investigate the confusion of pose and conformational heterogeneity, particular in the case of state population estimations by HRAs. In this dataset, synthetic particle images were generated from open and closed atomic structures, adapted from PDB IDs 6UIV and 6UIW, respectively. To ensure that the two structures had the same number of atoms, the ruthenium red molecule and 7 N-terminal residues were removed from the closed, 6UIW structure. Each structure was equilibrated in an explicit membrane via molecular dynamics simulations, such that the images were generated from a slightly different structure than the PDB. All simulations were set up using CHARMM-GUI (27) and simulations were performed with the GROMACS 2022 software package (28). The Charmm36m force-field with the TIP3P water model was used (29). The models were embedded in a POPC membrane for simulation with 150 mM NaCl plus neutralizing ions. Each system was energy minimized using steepest descent followed by a multi-step equilibration lasting a total of 375 ps. In the first two steps, the system was equilibrated in a canonical (NVT) ensemble with an integration time step of 1 fs for 50 ps each, maintaining a temperature of 310.15 K using the Berendsen thermostat (30). During this multi-step equilibration, position restraints were applied on lipid and protein heavy atoms, starting at 1,000 and 4,000 kJ/(mol/nm2), respectively, and then gradually lowered in every step until reaching zero in the final step. During the equilibration steps, we established a pressure of 1 bar using the Berendsen barostat with a characteristic time of 1 ps (30). Images were generated directly from non-hydrogen atomic coordinates of these equilibrated structures using Gaussian mixture projection.

To generate synthetic images using CryoJAX (version 5), we first made STAR files using an exact conformational open:closed ratio of 0.185:0.815 (5550 : 24450). We chose this ratio since it was far from uniform, while still included a few thousand images in the less populated state. We sampled non-uniform poses with zero-shift by first sampling unit-quaternions and converting them to Euler angles via `cryojax.simulator.EulerAnglePose.from_rotation_and_translation`. Rotational quaternions were sampled with `numpyro.distributions.ProjectedNormal` distribution, centered at the side view pose, with a concentration of 20, corresponding to an isotropic deviation of 9 +/- 4 deg in geodesic distance. A small fraction of uniform poses of 3 percent (900 particles) were drawn from a uniform rotational distribution, with the remaining 97 percent (29300) non-uniform. These numbers were chosen after testing single rounds of 2D and 3D classification at various SNR ratios, with the goal of making the task an appropriate level of difficulty for a community challenge.

The exact microscope parameters were supplied in the STAR file. The CTF defocus was uniformly drawn in a range of 1-2 um, with no astigmatism. We used other typical microscope and image formation parameters (300 keV beam voltage, 2.7 mm spherical aberration, 0.1 amplitude contrast, 1.2 Å pixel size, 256 pixel box size). We used Gaussian white noise (`cryojax.simulator.UncorrelatedGaussianNoiseModel(...,variance=1,signal_scale_factor=...)`, where `signal_scale_factor` was log-uniform sampled from 0.001 to 0.01. The specimen density was generated from non hydrogen protein PDB atom coordinates using `cryojax.simulator.GaussianMixtureProjection` with `cryojax.simulator.PengScatteringFactorParameters` scattering parameters, which generates 2D image density directly from atoms. Finally, we masked the image with `cryojax.ndimage.transforms.CircularCosineMask` using a radius of 128 and rolloff width of 0. Listing 1 shows a code snippet.

```python
import cryojax.simulator as cxs
image_config = cxs.BasicImageConfig(
        shape=(256, 256),
        pixel_size=1.2,
        voltage_in_kilovolts=300,
    )

```

```
transfer_theory = cxs.ContrastTransferTheory(
        ctf=cxs.AstigmaticCTF(
            defocus_in_angstroms=...,
            astigmatism_in_angstroms=0,
            astigmatism_angle=0,
            spherical_aberration_in_mm=2.7,
        ),
        amplitude_contrast_ratio=0.1,
        phase_shift=0,
    )

mask = tf.CircularCosineMask(
        coordinate_grid=...,
        radius=128,
        rolloff_width=0,
    )
signal_region = mask.array == 1

snr = ...

image_model = cxs.make_image_model(
        volume_parametrization=...,
        image_config=image_config,
        pose=...,
        transfer_theory=transfer_theory,
        volume_integrator=cxs.GaussianMixtureProjection(),
        normalizes_signal=True,
        signal_region=signal_region,
    )
noise_model = cxs.UncorrelatedGaussianNoiseModel(
        image_model,
        variance=1.0,
        signal_scale_factor=jnp.sqrt(snr),
    )
noise_model.sample(...)
```
Listing 1 | CryoJAX code snippet

[2] Alister Burt, Bogdan Toader, Rangana Warshamanage, Andriko Von Kügelgen, Euan Pyle, Jasenko Zivanov, Dari Kimanius, Tanmay A. M. Bharat, and Sjors H. W. Scheres. An image processing pipeline for electron cryo-tomography in <span style="font-variant:small-caps;">RELION</span> -5. 14(11):1788–1804. ISSN 2211-5463, 2211-5463. doi: 10.1002/2211-5463.13873. URL https://febs.onlinelibrary.wiley.com/doi/10.1002/2211-5463.13873.

[3] Ali Punjani, John L Rubinstein, David J Fleet, and Marcus A Brubaker. cryoSPARC: algorithms for rapid unsupervised cryo-EM structure determination. 14(3):290–296. ISSN 1548-7091, 1548-7105. doi: 10.1038/nmeth.4169. URL https://www.nature.com/articles/nmeth.4169.

[10] John Jumper, Richard Evans, Alexander Pritzel, Tim Green, Michael Figurnov, Olaf Ronneberger, Kathryn Tunyasuvunakool, Russ Bates, Augustin Žídek, Anna Potapenko, Alex Bridgland, Clemens Meyer, Simon A. A. Kohl, Andrew J. Ballard, Andrew Cowie, Bernardino Romera-Paredes, Stanislav Nikolov, Rishub Jain, Jonas Adler, Trevor Back, Stig Petersen, David Reiman, Ellen Clancy, Michal Zielinski, Martin Steinegger, Michalina Pacholska, Tamas Berghammer, Sebastian Bodenstein, David Silver, Oriol Vinyals, Andrew W. Senior, Koray Kavukcuoglu, Pushmeet Kohli, and Demis Hassabis. Highly accurate protein structure prediction with alphafold. *Nature*, 596(7873):583–589, 2021. ISSN 1476-4687. doi: 10.1038/s41586-021-03819-2. URL http://dx.doi.org/10.1038/s41586-021-03819-2.

[11] Keitaro Yamashita, Colin M. Palmer, Tom Burnley, and Garib N. Murshudov. Cryo-em single-particle structure refinement and map calculation using servalcat. *Acta Crystallographica Section D Structural Biology*, 77(10):1282–1291, 2021. ISSN 2059-7983. doi: 10.1107/s2059798321009475. URL http://dx.doi.org/10.1107/S2059798321009475.

[12] Jas Kalayan, James Gebbie-Rayet, and Harry Swift. aiida-gromacs, 2026. URL https://zenodo.org/doi/10.5281/zenodo.18838601.

[13] Jas Kalayan, James Gebbie-Rayet, and Harry Swift. aiida-amber, 2026. URL https://zenodo.org/doi/10.5281/zenodo.18835330.

[14] Elizabeth Jurrus, Dave Engel, Keith Star, Kyle Monson, Juan Brandi, Lisa E. Felberg, David H. Brookes, Leighton Wilson, Jiahui Chen, Karina Liles, Minju Chun, Peter Li, David W. Gohara, Todd Dolinsky, Robert Konecny, David R. Koes, Jens Erik Nielsen, Teresa Head-Gordon, Weihua Geng, Robert Krasny, Guo-Wei Wei, Michael J. Holst, J. Andrew McCammon, and Nathan A. Baker. Improvements to the apbs biomolecular solvation software suite. *Protein Science*, 27

(1):112–128, 2018. doi: https://doi.org/10.1002/pro.3280. URL https://onlinelibrary.wiley.com/doi/abs/10.1002/pro.3280.

[15] David A. Case, Hasan Metin Aktulga, Kellon Belfon, David S. Cerutti, G. Andres Cisneros, Vinicius Wilian D. Cruzeiro, Negin Forouzesh, Timothy J. Giese, Andreas W. Gotz, Holger Gohlke, Saeed Izadi, Koushik Kasavajhala, Mehmet C. Kaymak, Edward King, Tom Kurtzman, Tai-Sung Lee, Pengfei Li, Jian Liu, Tyler Luchko, Ray Luo, Madushanka Manathunga, Matias R. Machado, Hai Minh Nguyen, Kurt A. O'Hearn, Alexey V. Onufriev, Feng Pan, Sergio Pantano, Ruxi Qi, Ali Rahnamoun, Ali Risheh, Stephan Schott-Verdugo, Akhil Shajan, Jason Swails, Junmei Wang, Haixin Wei, Xiongwu Wu, Yongxian Wu, Shi Zhang, Shiji Zhao, Qiang Zhu, III Cheatham, Thomas E., Daniel R. Roe, Adrian Roitberg, Carlos Simmerling, Darrin M. York, Maria C. Nagan, and Jr. Merz, Kenneth M. Ambertools. *Journal of Chemical Information and Modeling*, 63(20):6183–6191, 10 2023. ISSN 1549-9596. doi: 10.1021/acs.jcim.3c01153. URL https://doi.org/10.1021/acs.jcim.3c01153.

[16] David A. Case, David S. Cerutti, Vinicius Wilian D. Cruzeiro, Thomas A. Darden, Robert E. Duke, Mahdieh Ghazimirsaeed, George M. Giambasu, Timothy J. Giese, Andreas W. Gotz, Julie A. Harris, Koushik Kasavajhala, Tai-Sung Lee, Zhen Li, Charles Lin, Jian Liu, Yinglong Miao, Romelia Salomon-Ferrrer, Jana Shen, Ryan Snyder, Jason Swails, Ross C. Walker, Jinan Wang, Xiongwu Wu, Jinzhe Zeng, Thomas E. Cheatham III, Daniel R. Roe, Adrian Roitberg, Carlos Simmerling, Darrin M. York, Maria C. Nagan, and Jr. Merz, Kenneth M. Recent developments in amber biomolecular simulations. *Journal of Chemical Information and Modeling*, 65(15): 7835–7843, 07 2025. ISSN 1549-9596. doi: 10.1021/acs.jcim.5c01063. URL https://doi.org/10.1021/acs.jcim.5c01063.
